# Characterization of the novel clinical isolate X-4 containing a new *tp0548* sequence-type

**DOI:** 10.1101/2019.12.16.877886

**Authors:** Dan Liu, Man-Li Tong, Li-Li Liu, Li-Rong Lin, Yu Lin, Tian-Ci Yang

**Author notes:** Corresponding author: Yu Lin and Tian-Ci Yang E-mail address and. These authors contributed equally to this work (Dan Liu, Man-Li Tong).

## Abstract

**Background:** A novel *tp0548* sequence-type was identified in one clinical isolate (X-4) from a patient diagnosed with primary syphilis in Xiamen, China. To precisely define and characterize this new clinical isolate, we performed further genome-scale molecular analysis.

**Methodology/Principal findings:** The alignment of all published *tp0548* genotypes revealed that this new genotype had a unique nucleotide substitution G->T at position 167, and the letter “ao” was assigned to the genotype. Phylogenetic analysis showed that the “ao” genotype belonged to the SS14-like clade of *Treponema pallidum* (TPA) strains. The genome of the X-4 isolate was then sequenced and analyzed, and the result of a multi-locus sequence analysis using a set of nine chromosomal loci showed that the X-4 isolate was clustered with a monophyletic group of TPA strains, which clearly identified the isolate as a TPA strain. Whole-genome phylogenetic analysis was subsequently conducted to corroborate the TPA strain classification of the X-4 isolate. And the isolate was genetically related to the SS14 strain, with 42 single nucleotide variations and 12 insertions/deletions. In addition, high intrastrain heterogeneity in the length of the poly G/C tracts was found in the TPAChi_0347 locus, which indicated that this gene was most likely involved in phase variation events. The first investigation of the length heterogeneity of the poly A/T tracts showed the variability of the ploy A/T was lower, and all the observed intrastrain variations fell within coding regions.

**Conclusions/Significance:** The study demonstrated the X-4 isolate was a TPA isolate containing a novel tp0548 sequence-type. The identification of intrastrain genetic heterogeneity at poly G/C tracts and poly A/T tracts of the isolate could provide a snapshot of the genes that potentially involved in genotype-phenotype variations. These findings provide an unequivocal characterization for better understanding the molecular variation of this emerging isolate.

**Author summary:** Three subspecies of *Treponema pallidum* (pallidum, pertenue, and endemicum) are increasingly showing overlap in terms of transmission and clinical manifestations. We recently identified a novel *tp0548* genotype in the X-4 isolate, which was obtained from an adult male with genital lesions. The novel genotype contained a unique nucleotide substitution G->T at position 167 and belonged to the SS14-like clade of TPA strains, as determined by phylogenetic analysis. We conducted an in-depth exploration of the genome of the X-4 isolate using the pooled segment genome sequencing method followed by Illumina sequencing. Multi-locus sequence analysis of nine chromosomal loci demonstrated that the X-4 isolate was clustered within a monophyletic group of TPA strains, which identified the isolate as a TPA strain. Whole-genome phylogenetic analysis subsequently corroborated the TPA strain classification of the X-4 isolate and revealed that the isolate was very closely related to the SS14 strain, with 42 single nucleotide variations and 12 insertions/deletions. In addition, characterization of the intrastrain heterogeneity in the lengths of homopolymeric tracts in the X-4 isolate showed that the heterogeneity of the poly G/C tracts was greater than that of the poly A/T tracts, and high poly G/C tract diversity was observed in the TPAChi_0347 locus.

## Introduction

Human pathogenic treponemas include three subspecies of *Treponema pallidum*, namely, *T. pallidum* subsp. *pallidum* (TPA), *T. pallidum* subsp. *pertenue* (TPE) and *T. pallidum* subsp. *endemicum* (TEN), and these subspecies are the causative agents of venereal syphilis (syphilis), yaws and endemic syphilis (bejel), respectively. These three subspecies share more than 99.5% sequence identity with each other and are indistinguishable by morphology and serology alone [1–3].

The oft-stated belief was that these three subspecies could be distinguished based on their mode of transmission, clinical manifestations and host specificity, but this probably appears to not be the case. In a previous study, one Paris isolate 11q/j was obtained from a syphilis-like primary genital lesion and was first reported as a syphilis case containing a novel *tp0548* sequence-type “j” by enhanced CDC (ECDC) typing system [4]. However, the novel *tp0548* genotype “j” was further found to be similar to those of TPE strains, and the “q” RFLP pattern of the *tpr* subfamily II genes was also consistent with that of most TPE strains. Hence, this 11q/j isolate was defined as an imported case of yaws [5]. After extensive molecular locus analysis, the isolate was finally clearly characterized as a TEN isolate that contained two recombination events in the *tp0548* and *tp0488* loci, resulting in sequences similar to those of TPE and TPA strains, respectively [6, 7]. Moreover, Noda *et al*. [8] performed sequencing-based molecular typing analysis of 92 Cuba samples from patients diagnosed with syphilis based on clinical and epidemiological data and found the infectious agent of nine samples was actually classified as TEN. Similarly, TEN has recently been found to spread among men who have sex with men (MSM) in Japan [9, 10]. It reminds us of the need to carefully characterize any isolate with a novel or unusual molecular sequence-type [8].

Recently we identified a novel *tp0548* genotype of an isolate which was obtained from a patient diagnosed with syphilis based on clinical manifestation and serology results. The ECDC subtype of the isolate was 13/d/? and the profile by the new proposed multilocus sequencing typing (MLST) scheme [11] was 1.?.8. After careful consideration, we performed in-depth molecular analysis at the genome scale to determine whether the infectious agent of this isolate was a novel TPA strain or another treponemal subspecies that was clinically misdiagnosed. Importantly, we further investigated the genome sequence to get insight into the genetic variation of this novel isolate.

## Materials and Methods

### Ethics statement

This study was approved by the Ethics Committee of Zhongshan Hospital, Xiamen University, and conformed with the Declaration of Helsinki. Written informed consent was obtained according to institutional guidelines prior to the study. All rabbit experiments strictly followed the parameters outlined by the Institutional Animal Care and Use Committee (IACUC) and were approved by the animal experimental ethics committee of School of Medicine, Xiamen University.

### Source of the X-4 isolate and DNA extraction

Ulcer swab samples were collected from a 42-year-old male patient who attended the Outpatient Department of Zhongshan Hospital, Xiamen University, for indurated painful genital ulceration. Syphilis serology showed positivity for *T. pallidum* particle agglutination (TPPA) and reactive plasma reagin (RPR) at 1:32. Samples of the ulceration that yielded positive results through dark field microscopy were immediately propagated in New Zealand White rabbits by intratesticular inoculation for enriching the isolate as previously published method [12].

DNA from the obtained treponemal suspensions was extracted using the QIAamp DNA mini kit (Qiagen Inc., Chatsworth, CA, USA), and all procedures were conducted strictly following the manufacturer’s instructions [13]. The extracted DNA was then positively determined by qPCR targeting *tp0574* [14]. The sequencing samples used in this study were obtained from the third passage in rabbits.

### Whole-genome sequencing and *de novo* assembly of the X-4 genome

The genome of the X-4 isolate was determined using the pooled segment genome sequencing (PSGS) method as reported previously [15, 16]. Library construction and sequencing were performed by Beijing Novogene Bioinformatics Technology Co., Ltd., with an Illumina HiSeq 1500 platform in the paired-end mode. Prior to genome assembly, quality control evaluation on the raw sequencing data using the Trimmomatic tool was performed. The Illumina sequencing reads corresponding to individual pools were handled separately and *de novo* assembled using SOAP *de novo*. The resulting contigs from the X-4 isolate were aligned to sequences from the *T. pallidum* subsp. *pallidum* Nichols strain using Lasergene software (DNASTAR, Madison, WI, USA).

All genome gaps and discrepancies were then filled and resolved by Sanger sequencing to obtain the complete genome sequence of the X-4 isolate. To determine the number of repetitions within the *arp* (*tp0433*) gene, the gene was amplified and sequenced using the primers (F: 5’-GTCGTTACCCGTTGTATTGC-3’ and R: 5’-CCTTCCCTTCCGTTCCTT-3’). Also, the size of the amplicon was determined by comparison with a 2-kb DNA marker with the use of Quantity One software (version 4.6.7; Biorad), and the number of repetitions was estimated by comparing the size of the product amplified from the Nichols strain, which contains 14 repetitions. The repetitive sequences within the *tp0470* gene were amplified and sequenced using the sense primers (5’-TCACGTCGTTCCGTCAGT-3’) and antisense primer (5’-GTAGGGTCCAAGCGAATAAG-3’). The complete genome was annotated using the NCBI PGAAP pipeline and the genes were tagged with FFV11_prefixes.

### Evaluation of the classification of the X-4 isolate based on the *tp0548* locus and nine chromosomal loci

BioEdit Sequence Alignment software v7.2.5 (Ibis Biosciences, Carlsbad, CA, USA) was employed to compare an 83-bp region of the *tp0548* locus [17] from the X-4 isolate with all available data in GenBank. Nine chromosomal loci (i.e., 16S rDNA, *tp0136, tp0326, tp0367, tp0488, tp0548, tp0859, tp0861* and *tp0865*) were analyzed using a method (multi-locus sequence analysis, MLSA) proposed by Noda *et al*. [8]. Sequence Matrix 1.8 software was used for sequence concatenation. The phylogenetic tree was generated with the Maximum Likelihood method in the Tamura-Nei model.

### Whole-genome comparisons

Whole-genome-based phylogenetic tree of the X-4 isolate and six other TPA strains, including the Nichols (CP004010.2), SS14 (CP004011.1), Mexico A (CP003064.1), DAL-1 (CP003115.1), Chicago (CP001752.1) and Sea 81-4 (CP003679.1) strains, was constructed with the Maximum Likelihood method. Due to the high intrastrain diversity in the *tprK* gene and the presence of repetitive sequences in the *arp* gene, those two genes were excluded from the analysis [18]. We then performed a genomic comparison of the X-4 isolate against its most closely related ancestral strain using the MUMmer and LASTZ tools. Nucleotide positions located within homopolymeric tracts (defined as a stretch of seven or more identical nucleotides) were excluded from the analysis [19].

### Evaluation of intrastrain heterogeneous G/C and A/T regions in DNA homopolymeric tracts of the X-4 isolate

Based on a previously published principle [18], an in-house Perl script was developed and used for the specific extraction of DNA sequences to determine the intrastrain sequence length variability in DNA homopolymeric tracts of guanosine/cytosine (G/C regions) and adenine/thymine (A/T regions). The exact number of different base counts in the tracts was determined, and the precise relative frequencies of different base counts within the intrastrain variable tracts were calculated. To ensure valid results with respect to the variability or conservation of homopolymeric tracts, the analysis quality was improved according to previous described [18]. Only homopolymeric tracts with a length greater than seven bases were included. The prevailing length of each G/C and A/T tract was used in the final genomic sequence.

### Accession numbers

The complete genome sequences of the X-4 isolate were deposited in GenBank under accession number CP040555. The raw data sequences of the X-4 isolate obtained in this study were deposited in the SRA database (BioProject ID: PRJNA544173) under the following BioSample accession number: SAMN11812273.

## Results

### Whole genome sequencing of the X-4 isolate and *de novo* assembly of the genomes

The X-4 isolate genome was sequenced using the PSGS approach. Illumina sequencing of the X-4 isolate genome yielded 39,782,973 paired reads and 4,800,380,928 total bases, with an average coverage depth of 4201×. A total of 4, 13, 4, and 8 contigs for pools 1–4 of the X-4 isolate were obtained by *de novo* assembly. The detailed characteristics of the Illumina sequencing and *de novo* assembly are shown in S1 Table. We separately aligned the resulting contigs to the Nichols genome, and 13 genome gaps and discrepant regions were filled and resolved by Sanger sequencing. The consensus sequences for the individual pools were then used to compile the genome sequence, and the results yielded the complete circular genome of the X-4 isolate, which contained 1,139,838 bp (S1 Fig).

### Classification of the X-4 isolate

#### Analysis of the X-4 isolate based on the *tp0548* locus

An analysis of the X-4 isolate *tp0548* locus (nucleotides 130-212 according to *tp0548* in the Nichols strain genome [AE 000520.1]) with published data, including a known TPA pattern and a known TPE pattern *in silico*, showed that the *tp0548* genotype in the X-4 isolate was novel. The known *tp0548* genotypes (a-ak) were assigned in a unique list containing TPA strains and TPE strains [20–22], and three other *tp0548* genotypes of TPA strains were recently published: two were defined as type “r” and “s” by Grillová *el al*. [11], and one was defined as type “y” by Kumar *et al*. [23]. For consistency with the already extensive list of *tp0548* genotype sequences (a-ak), we redefined types “r”, “s” and “y” as “al”, “am” and “an” and added the new *tp0548* sequence type of the X-4 isolate to the updated list as type “ao” (Fig. 1). Phylogenetic analysis of all *tp0548* genotypes divided the treponemas into three clades (S2 Fig) and the novel genotype “ao” was found to belong to the SS14-like clade of TPA strains.

**Fig 1.**
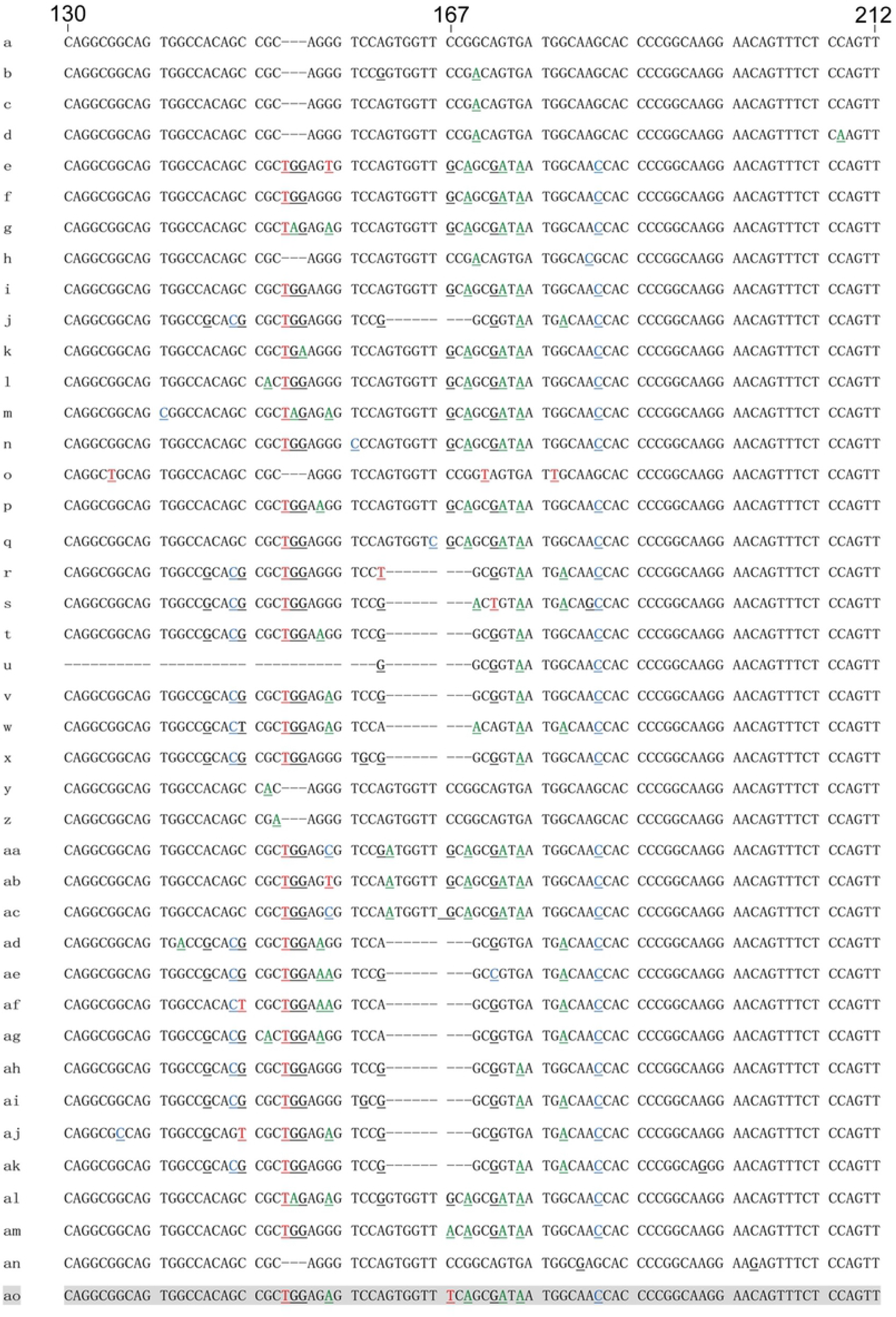
All identified *tp0548* genotypes by sequencing analysis of an 83-bp region (nucleotides 130-212 according to *tp0548* in the Nichols strain genome) The new genotype “ao” is shown in gray.

Because of the novel *tp0548* genotype, the ECDC subtype (13d/ao) of the isolate was a new strain type. Comparing the predominant subtype (14d/f) in clinical isolates of China, the *tp0548* genotype of the X-4 isolate contains two nucleotide substitutions, which were G->A at position 55 and G->T at position 167 (cognate to the Nichols strain), respectively. The nucleotide substitution G->T at position 170 was unique when aligning with all the *tp0548* genotype (Fig. 1).

#### Analysis of the X-4 isolate by MLSA

To further classify the infectious agent of this isolate, we performed MLSA based on nine different chromosomal loci (16S rDNA, *tp0136, tp0326, tp0367, tp0488, tp0548, tp0859, tp0861* and *tp0865*) The Maximum Likelihood phylogenetic tree clearly divided the TPA, TPE and TEN strains into three monophyletic clades (Fig. 2), and the X-4 isolate formed part of a monophyletic group consisting of TPA strains (SS14 and PT_SIF0751).

**Fig 2.**
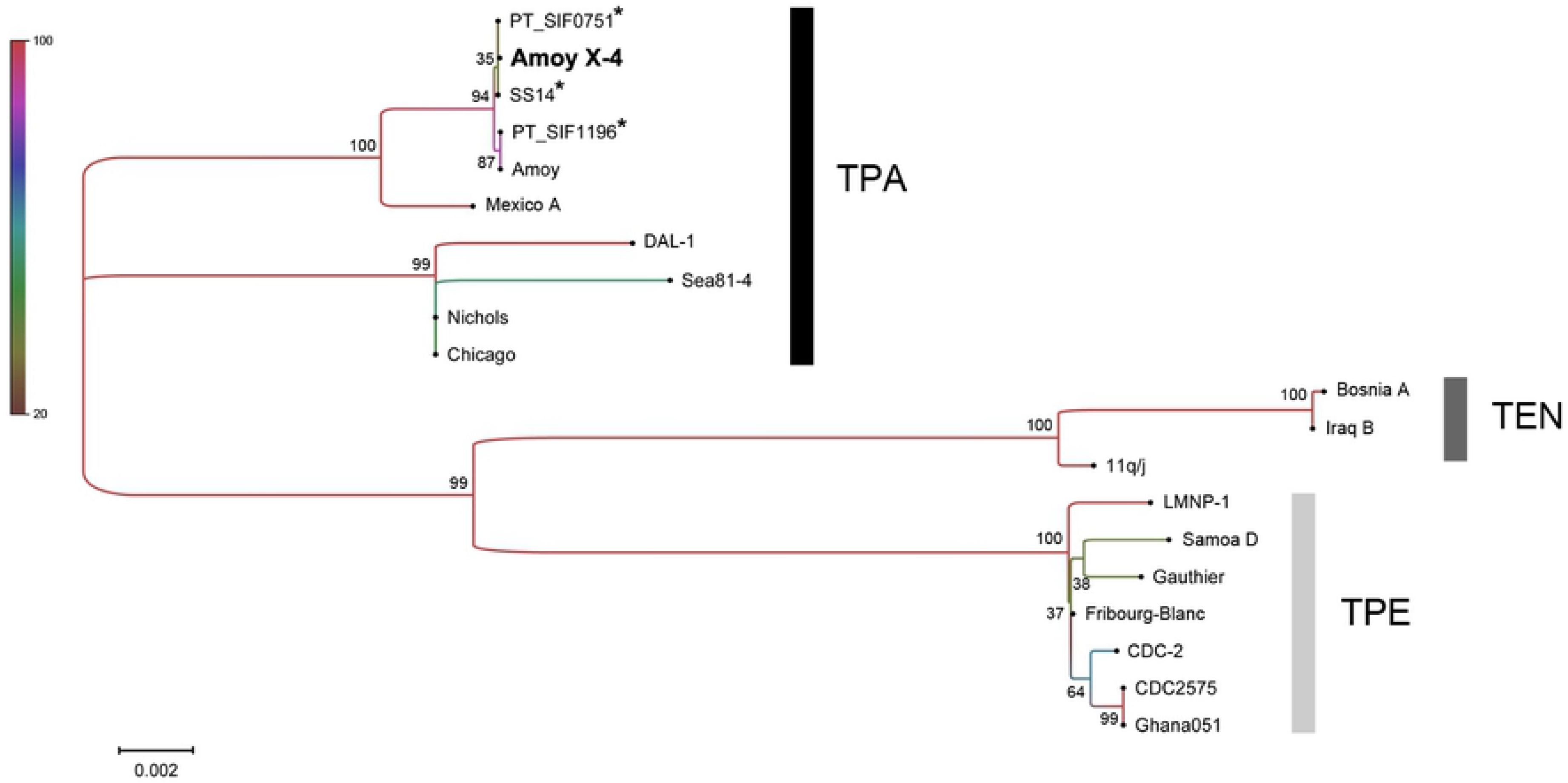
Analysis of the MLSA scheme based on all TPA/TPE/TEN strains with available sequences of nine loci. The phylogenetic tree was constructed using the Maximum Likelihood method and the Tamura-Nei model. The bootstrap support (out of 1000 replicates) is indicated. The asterisks indicate the existence of other strains with identical concatenated sequences of the nine tested loci[8].

Similarly, the profile (1.48.8) of the X-4 isolate by MLST was a newly identified profile. Comparing already identified allelic variant in *tp0548* locus, the *tp0548* locus of the X-4 isolate was showed to only contain two nucleotide substitutions at position 55 and at position 167, as above described.

#### Analysis of the X-4 isolate at the whole-genome scale

A whole-genome phylogenetic analysis of the X-4 isolate and six TPA reference strains available at GenBank was performed and the results showed that the seven TPA strains were grouped into three clades, and among these clades, the X-4 isolate was segregated in a separate branch and closely related to the SS14 strain (Fig. 3).

**Fig 3.**
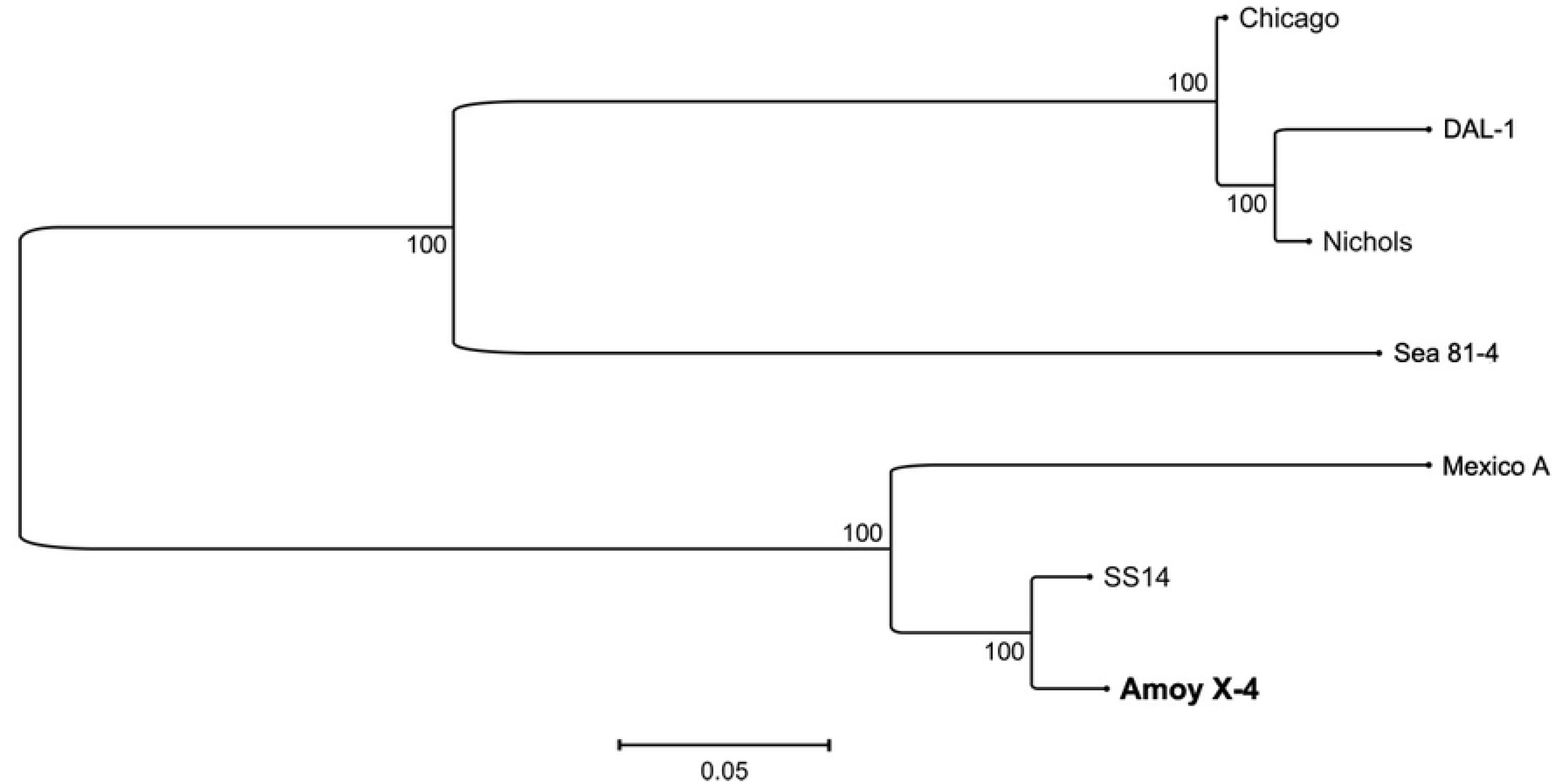
Whole-genome based classification of the X-4 isolate. Whole-genome-based Maximum Likelihood phylogenetic tree was constructed (the *tprK* and *arp* genes were excluded). The bootstrap values based on 1,000 replicates are shown next to the corresponding branch nodes.

We subsequently analyzed the genome of the X-4 isolate by referring to the SS14 genome with respect to the occurrence of nucleotide diversity. Overall, we found 42 variable nucleotide positions, 39 of these were located in coding regions, and 27 of these 39 positions were nonsynonymous variant sites (Fig. 4). Moreover, 12 insertions/deletions in genomic regions were identified in the X-4 genome relative to the SS14 genome, and five of these mutations were in coding regions (Table 1). Among the five mutations, four deletions caused frameshifts, resulting in omission of the annotations TPASS_RS00905, TPASS_RS05250, TPASS_RS04250 and TPASS_RS05155 according to the SS14 strain genome. Notably, a 9-nt-long insertion (TCCTCCCCC) at 1050766-1052319 (according to the SS14 genome) was found in a coding region. We further investigated the corresponding gene (annotated as the *tp0967* gene) and found that its genomic region had a different number of 9-nt-long (TCCTCCCCC) repetitive sequences. Specifically, the SS14 strain contained three of these repetitions, whereas the genome of the X-4 isolate contained four of these repetitive sequences. Besides, we also detected macrolide resistance-causing mutation in the X-4 strain. It revealed the isolate harbored the A2058G mutation as that presented in the SS14 strain.

**Fig 4.**
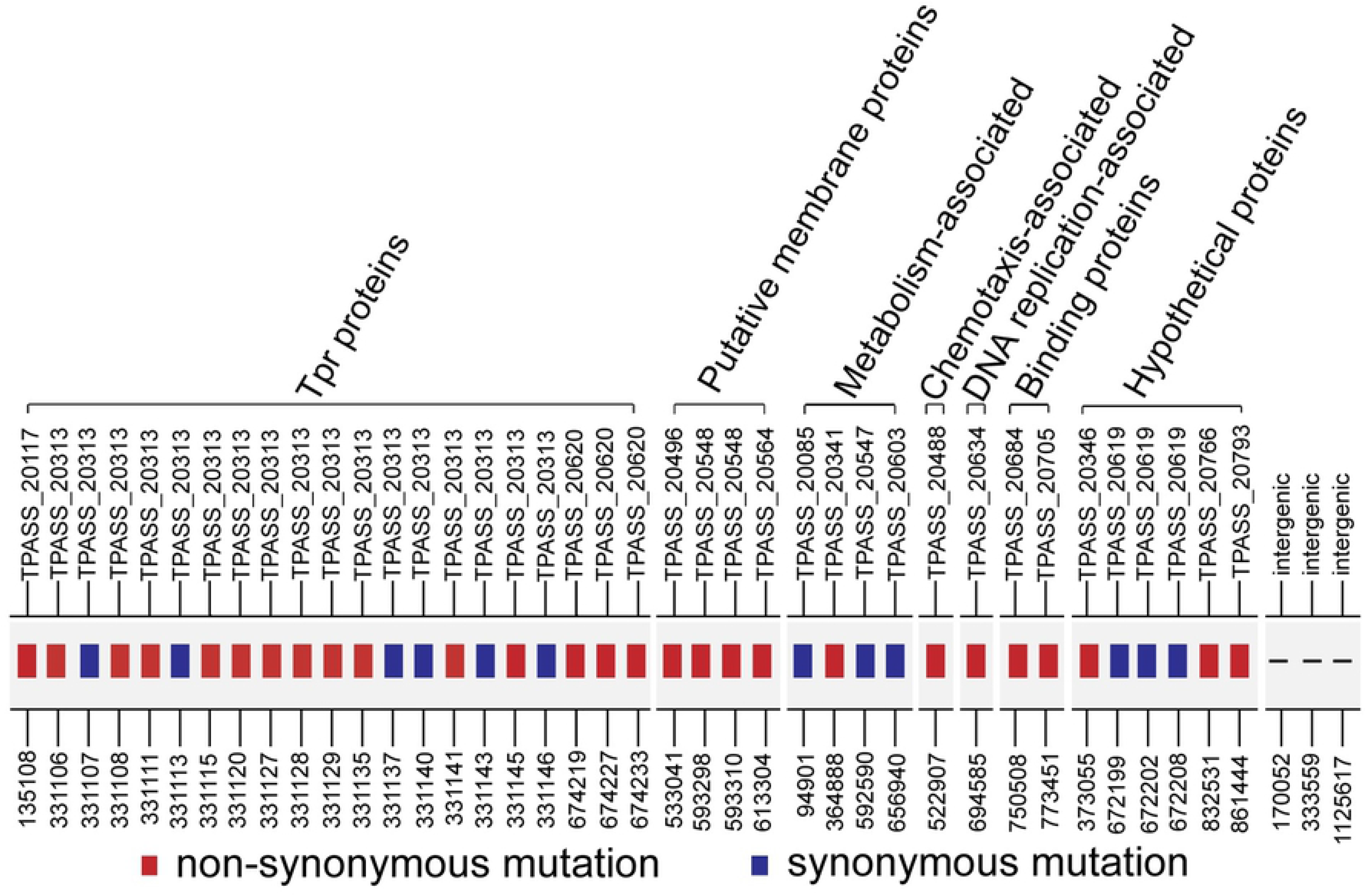
Mutations in the genes of the X-4 isolate compared with those of the SS14 strain

**Table 1.**
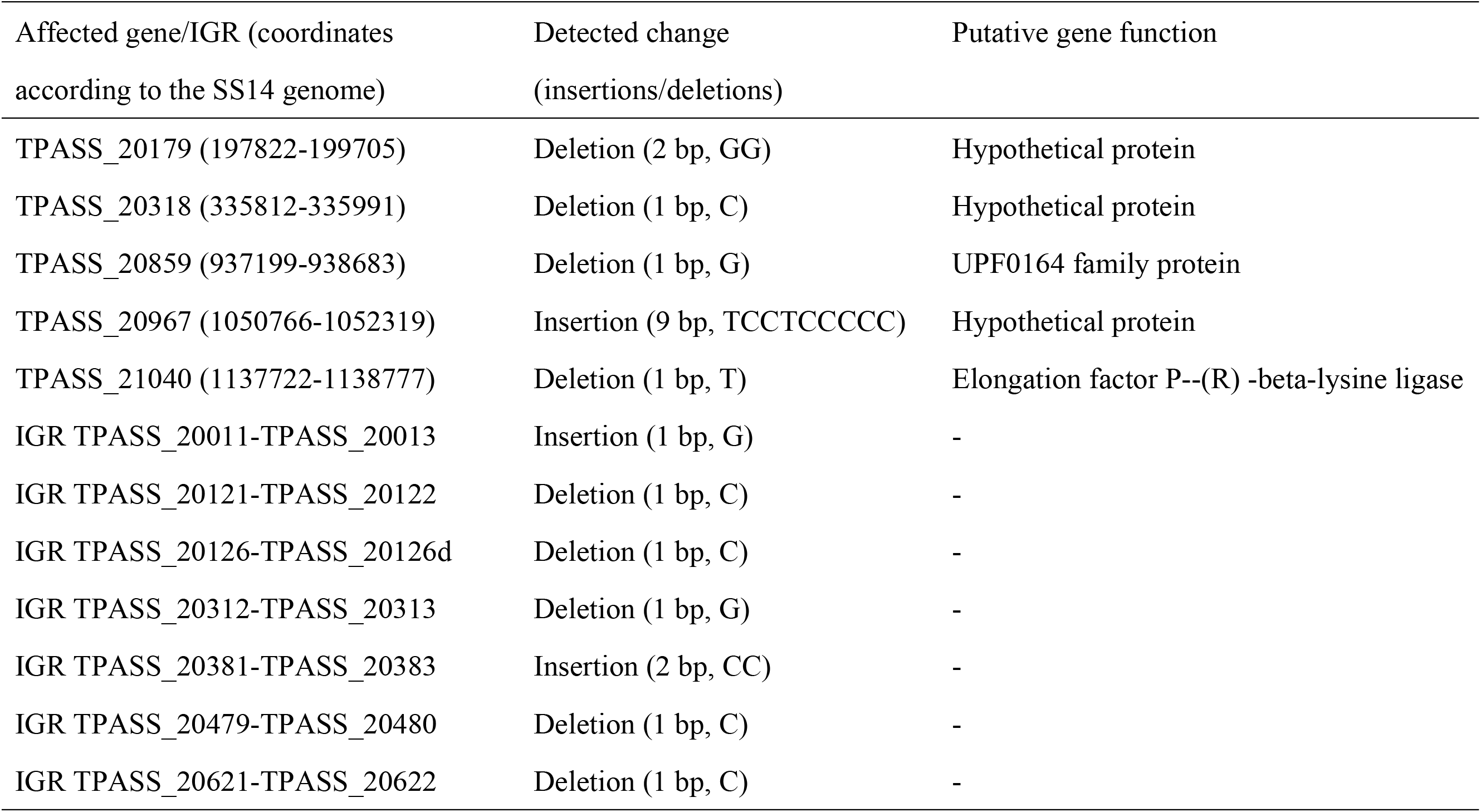
List of insertions/deletions in the X-4 isolate compared with the sequenced SS14 strain

### Special intrastrain genetic heterogeneity in homopolymeric tract lengths (G/C and A/T regions) of the X-4 isolate

Cumulative evidence demonstrates that genetic variability in homopolymeric tracts within an infectious population plays an important role in phenotypic variation. In this study, the genome of the X-4 isolate demonstrated remarkable intrastrain genetic heterogeneity at homopolymeric tracts. Altogether, 100 chromosome-dispersed poly G/C tracts were identified in the X-4 genome and analyzed. Among the 100 tracts, 30 of the poly G/C tracts showed intrastrain genetic variability and were equally located within predicted coding regions and regulatory regions (Fig. 5a and S2A Table). Notably, the frequencies of the dominant base counts (11 poly G/C counts) and the secondary base counts (12 poly G/C counts) in the TPAChi_0347 locus reached 39.5% and 34.9%, respectively. As such, the variable number of poly (G/C) tracts in TPANIC_0347 revealed up to five types of base count (Fig. 5a). Additionally, we noted that most of the heterogeneity in the putative regulatory regions was located upstream of genes belonging to the *tpr* family.

**Fig 5.**
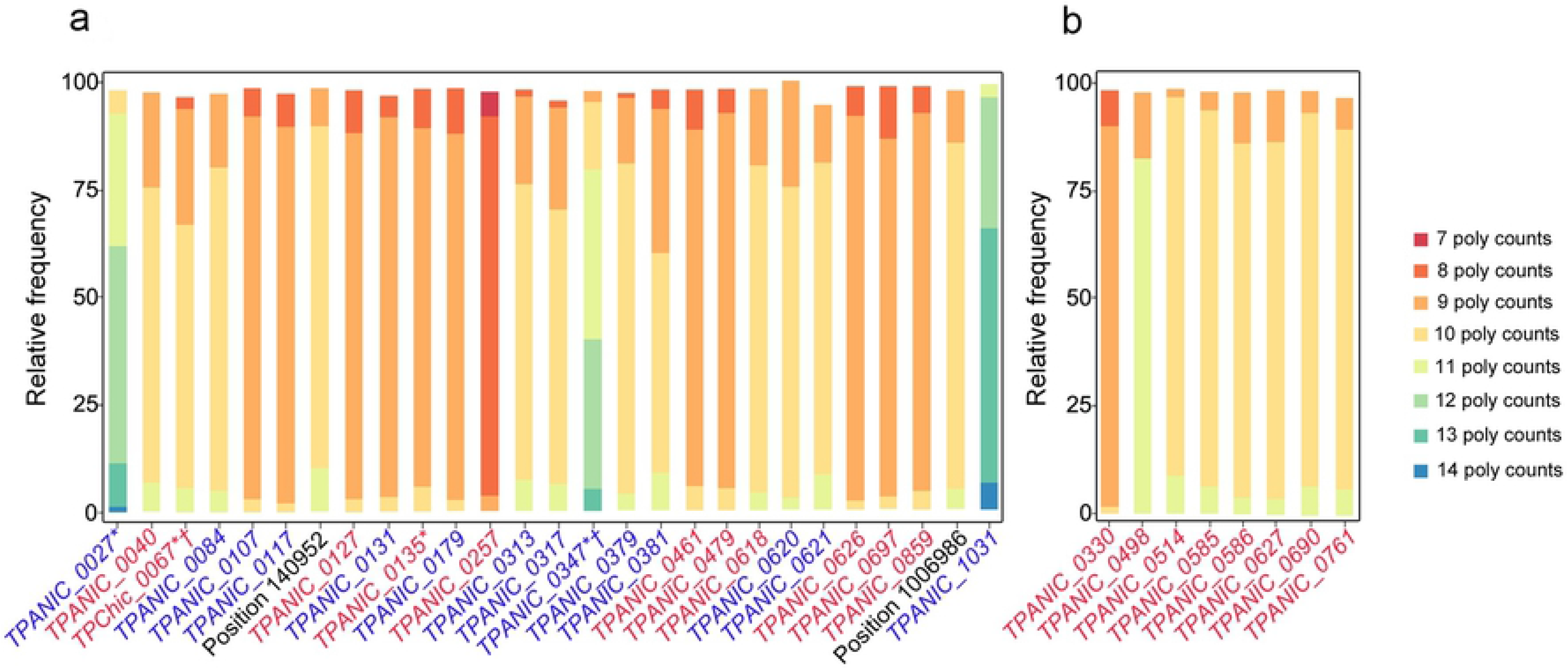
Genetic heterogeneity in the lengths of homopolymeric tracts (G/C and A/T regions) in the genome of the X-4 isolate. Each graph displays the relative percentage of sequence reads with a particular base count in the genome of the X-4 isolate. The genes displayed below the graphs are potentially affected by phase variation coordinating to the Nichols genome (CP004010.2) and are colored according to the relative position of the poly G/C tract in a putative regulatory region (blue), within a coding region (red) or in an unpredictable region (black) (presenting the Nichols genome position). Genetic heterogeneity in the lengths of a) poly G/C tracts and b) poly A/T tracts. * indicates incongruences regarding the annotation of the gene. † indicates no annotation in the Nichols genome, and thus, the gene refers to the genome of the Chicago strain (CP001752.1). The list of potential affected gene by poly G/C tracts referenced the genes identified in Pinto’s study.

Meanwhile, we investigated the variability in the lengths of poly A/T regions in the genome of the X-4 isolate. Although 557 chromosome-dispersed poly A/T tracts were identified, only eight tracts showed intrastrain variability. The relative frequencies of the dominant base counts of these eight homopolymeric tracts were almost 80% (Fig. 5b and S2B Table). Moreover, all eight tracts were distributed within coding regions associated with general metabolism, DNA repair and transport.

## Discussion

In the present study, we found that a *tp0548* sequence-type in the clinical isolate X-4 that had not been previously identified through the ECDC or MLST typing system [4, 11, 17, 20, 22–29]. Based on an analysis of an 83-bp region in the *tp0548* locus, the phylogenetic tree revealed that this novel sequence-type, “ao”, clustered with SS14-like genotypes. Due to the high similarity in the genome sequences and the possible occurrence of recombination events among the three human pathogenic treponemas (TPA/TPE/TEN strains) [7, 30], we decided to perform an in-depth molecular analysis to clearly classify and characterize this new isolate. Here, we used the PSGS approach, which has been successfully applied to amplify the genomes of *T. pallidum* subspecies [15, 16, 31], to analyze the genome sequence of the X-4 isolate.

As reported in a previous study [8], an analysis of nine chromosomal regions could clearly distinguish among TPA, TEN and TPE. Thus, we extracted the corresponding sequences from the assembled contigs of X-4 and conducted a phylogenetic analysis. The results were consistent with the classification of the X-4 isolate as a TPA strain. Moreover, whole-genome-based phylogenetic analysis indicated that the X-4 isolate and SS14 strain might have originated from the same ancestor, which further confirmed that the X-4 isolate was a TPA strain.

Comparative genomic results showed that the X-4 isolate had a 9-nt-long repetitive sequence inserted in the 1050766-1052319 region (in the hypothetical protein-encoding gene *tp0967*). Previous studies have demonstrated that this genomic region is variable in most pathogenic treponemal strains [1, 16]. We analyzed the available treponemal sequences in the NCBI database and found that the number of sequence repeats (TCCTCCCCC) varied from one to four in all the investigated strains. Genomic analyses have revealed that the genomes of the three subspecies are highly similar [1, 3]; thus, simple genetic differences that occurred during the evolution of these genomes might provide important evolutionary hints. In *Legionella pneumophila*, the number of repeats could reflect the strain origin [32]. Thus, the gain or loss of repeated sequences resulting in variability in the *tp0967* gene within the three subspecies might reflect the origin of a strain. In fact, only in TPA strains did we find a completely identical sequence pattern to that of the X-4 isolate (S3 Table). As novel TPA isolate, the X-4 isolate was closely related to the SS14 strain, which was consistent with previous phylogenetic studies on *T. pallidum* strain showing that currently most prevalent strains worldwide are SS14-like strains harboring macrolide resistance-causing mutation [33–35].

Additionally, the X-4 isolate demonstrated remarkable intrastrain genetic heterogeneity at homopolymeric tracts. The intrastrain heterogeneity of homopolymeric tracts within coding regions or putative regulatory regions has been found to be involved in phenotypic variation in many bacteria [36, 37], particularly bacteria with limited metabolic capabilities, e.g., *T. pallidum* [38, 39]. In this study, we found that most intrastrain in-length variable poly G/C tracts in the X-4 isolate were consistent with the results of a recent analysis of clinical syphilis isolates from Portugal [18], which indicates that these regions might show intrastrain variability in most TPA strains.

And most of the heterogeneity were found within the putative regulatory regions of the *tpr* family, which corroborated the finding by Giacani *et al*[38]. It is noteworthy that TPNIC_0126 was found to be transcriptionally affected by variations in the poly G/C tract length, which is consistent with regulation by phase variation [30]. These poly tracts in the X-4 genome were conserved with a 9-base count, which was similar to the findings obtained by Pinto et al [18]. In fact, we observed variations in the poly tracts of TPNIC_0126 in the X-4 isolate (eight and ten base counts), but their relative frequencies were low (3.5% and 4.5%, respectively). Additionally, theTPAChi_0347 locus in the X-4 isolate showed obvious diversity in the lengths of the homopolymeric tracts, which was similar to the findings obtained for the PT_SIF1252 strain [18]. Despite incongruences regarding the annotation of this gene [31, 40], similar results were obtained in the two studies, which suggests that the transcription or protein status (functional on/off) of the gene is more likely to be modulated by phase variation. Further experimental validation is necessary to confirm whether the gene is in the “on” or “off” state during *in vivo* growth of the X-4 isolate and to investigate whether its activity correlates with infection development.

Besides, we first analyzed the variability in the length of poly A/T tracts of the X-4 strain in this study. Compared with the poly G/C tracts, the poly A/T tracts in the genome of the X-4 isolate showed decreased diversity, and the relative frequency of the dominant base count was almost above 80%. No variable poly A/T tracts were found in the putative promoter regions, which suggests that the transcript levels of the genes in *T. pallidum* modulated by homopolymeric tracts might depend on poly G/C rather than poly A/T tracts. All variable poly A/T tracts were found within genes related to the pathogen’s general metabolism and DNA repair and transport, which indicates that heterogeneity in poly A/T tract might affect *T. pallidum*’s biology. As such, because the variability in the length of poly A/T tracts in human pathogenic treponemes has not been investigated, the heterogeneous poly A/T tracts found in this study could constitute a mapping of potential targets of phase variation by poly A/T tracts.

Finally, the limitations of our research should be discussed. First, the whole genome analysis was conducted only using the available complete genome sequence of TPA strains. Although the result clearly demonstrated that the X-4 strain was a TPA strain, it may be a relatively small number. Second, the results from the analysis of homopolymeric tracts in the X-4 isolate should be viewed with caution, particularly those obtained for tracts with low coverage.

This study investigated the X-4 isolate at genomic-scale to clearly characterize the isolate. The findings demonstrated the X-4 isolate was a TPA isolate containing a novel tp0548 sequence-type, which was closely related to the SS14 strain. Remarkable Intrastrain genetic heterogeneity at poly G/C tracts and poly A/T tracts found in the isolate could provide the information about the prioritizing loci involving genotype-phenotype variation.

## Acknowledgments

We thank prof. David Šmajs from Masaryk University for his guidance on using the PGSG method.

## Supporting information

**S1 Fig. Circular genome of the X-4 isolate** The outermost circle shows the genomic sequence position coordinates. From the outside to the center: Genes on the forward strand and on the reverse strand, functional annotation of the coding genes (based on the COG, KEGG and GO categories, respectively), RNA genes (tRNAs green, 5S RNAs dark brown, 16S rRNAs orange, 23S RNAs light brown), GC content, and GC skew.

**S2 Fig. Phylogenetic relationships of all *tp0548* genotypes** The phylogenetic tree was constructed using the Maximum Likelihood method with a bootstrap test of 1,000 replicates to evaluate the confidence. The bootstrap number (in percentage) is shown next to the corresponding node.

**S1 Table. Sequencing statistics for the genome of the X-4 isolate**

**S2 Table. Intrastrain genetic heterogeneity of poly G/C and A/T tracts in the genome of the X-4 isolate** A) Poly G/C tracts and B) poly A/T tracts in the genome of the X-4 isolate.

**S3 Table. Numbers of repetitive sequences (TCCTCCCCC) in the *tp0967* gene among treponemal strains**

